# EPAS1 regulates host antibacterial defense in response to bacterial quorum sensing

**DOI:** 10.1101/2025.10.09.681351

**Authors:** Heng Li, Xiaohan Kong, Binbin Cui, Xiaohui Li, Xia Li, Mingfang Wang, Liu-En Wang, Xiayu Chen, Wei Qian, Lian-Hui Zhang, Yongliang Zhang, Yinyue Deng

## Abstract

Hypoxia-inducible factors (HIFs) are a family of transcription factors that function as sensors of hypoxia. However, it remains unclear whether and how HIFs modulate host immunity by responding to other ligands. Here, we describe an immune regulatory function of HIF-2α (also known as EPAS1) in response to *cis*-2-dodecenoic acid (BDSF), a quorum sensing (QS) signal, secreted by *Burkholderia cenocepacia*. Upon ligand binding, EPAS1 activation leads to the production of inflammatory cytokines and the initiation of antibacterial defense. Inoculation with diffusible BDSF or liposome-encapsulated BDSF improves survival in mice infected with *B. cenocepacia* or *Acinetobacter baumannii* by preconditioning the immune system. Our findings reveal a previously unidentified role of EPAS1 as an intracellular pattern recognition receptor (PRR) for bacterial QS signal to control host antibacterial defense.

## Introduction

Hypoxia-inducible factors (HIFs) are central regulators of cellular adaptation to oxygen deprivation (*1, 2*). HIF proteins act as heterodimers composed of one α subunit, such as HIF-1α, HIF-2α (also known as endothelial PAS domain-containing protein 1, EPAS1), or HIF-3α, and one β subunit known as ARNT (aryl hydrocarbon receptor nuclear translocator), which together regulate transcriptional programs in response to hypoxia (*3-5*). Among the HIF family members, HIF-1α is broadly expressed and primarily mediates acute hypoxic responses, regulating the genes involved in glycolysis, angiogenesis, and cell proliferation (*6*). HIF-3α is thought to function primarily as a negative regulator of hypoxia-inducible signaling, modulating the activity of HIF-1α and EPAS1 (*7*). EPAS1 shows a more selective expression pattern in a cell context-dependent manner and is enriched in endothelial, epithelial, and immune cells, where it uniquely regulates erythropoiesis, vascular development, and inflammatory gene expression (*8-11*). It was recently indicated that gut microbe-derived metabolites affected EPAS1-mediated physiology (*12, 13*). However, it is unclear whether EPAS1 could sense the metabolites derived from pathogenic bacteria to modulate host immunity.

Quorum sensing (QS) is a process of cell-cell communication that coordinates bacterial group behaviors in a cell density-dependent manner (*14*). Diverse QS signals, including N-acyl homoserine lactones (AHLs) and diffusible signal factor (DSF) family signals, have been identified to regulate virulence, antibiotic resistance, biofilm formation, and adaptation to diverse environments (*15-19*). Notably, the QS molecule *cis*-2-dodecenoic acid (BDSF), originally identified from *Burkholderia cenocepacia*, modulates a wide range of bacterial phenotypes (*20-22*). Beyond intra- and interspecies signaling, QS molecules have emerged as critical mediators of cross-kingdom communications, directly interacting with host cell receptors, the aryl hydrocarbon receptor (AHR), to influence signaling transduction and immune responses (*23*). Thus far, it remains unknown whether QS signals can also mediate innate immune responses through other pattern recognition receptors (PRRs).

In this study, we demonstrate that EPAS1 functions as an intracellular PRR by binding and sensing the bacterial QS signal BDSF, which may represent a previously unrecognized pathogen-associated molecular pattern (PAMP) that regulates host immune response against bacterial pathogens. Intriguingly, we found that inoculation of BDSF QS signal protects mice against bacterial pathogens, regardless of whether the pathogens produce BDSF, suggesting that the EPAS1-dependent signaling axis elicits a moderate yet protective innate immune response. Together, our findings uncover a previously unrecognized host-pathogen communication axis mediated by EPAS1 and QS signals.

## Results

### BDSF from *B. cenocepacia* elicits immunostimulatory responses *in vitro*

To explore whether there are unidentified bacteria-derived small molecules that can regulate host immune responses, we treated murine RAW264.7 macrophages with the ethyl acetate extract of *B. cenocepacia* culture supernatant (BCS) and examined changes in cellular morphology, inflammatory gene expression, and cytokine production in the macrophage cells, using lipopolysaccharide (LPS) as a positive control. As shown in fig. S1A, treatment with LPS or *B. cenocepacia* H111 bacteria induced marked morphological alterations characterized by increased cell clustering and membrane irregularities, both indicative of macrophage activation. Notably, exposure to the extract of BCS also provoked morphological changes consistent with the activated macrophage phenotypes (fig. S1A). Quantitative RT-PCR analysis further demonstrated significant upregulation of *Tnfa* and *Il6* transcripts following treatment with LPS and H111 bacteria, while BCS elicited a moderate yet consistently detectable induction of these inflammatory genes, compared to the untreated cells (fig. S1B). Consistent with these transcriptional changes, elevated secretion of TNFα and IL-6 proteins in all treatment groups, with the strongest induction observed in response to LPS and H111 bacterial cells, and intermediate responses to BCS was observed (fig. S1C). Collectively, these results indicate that both bacteria and their secreted small molecules can activate macrophages and promote inflammatory responses.

To evaluate whether the defined QS molecules from the BCS can directly activate macrophages, we assessed the effects of C6-AHL (N-hexanoyl-L-homoserine lactone), C8-AHL (N-octanoyl-L-homoserine lactone), and BDSF, which are the QS signals produced by this pathogen (*16, 20*), on RAW264.7 cells. BDSF induced moderate morphological changes, whereas C6-AHL and C8-AHL exerted only weaker effects (Fig. 1A). Consistent with these observations, BDSF significantly upregulated *Tnfa* and *Il6* transcripts, whereas C6-AHL and C8-AHL elicited only modest changes in the expression of these genes (Fig. 1B). Consistent with the transcript data, BDSF induced higher secretion of TNFα and IL-6 compared to control and AHLs-treated groups, although these responses are still lower than those elicited by LPS (Fig. 1C). Together, these results indicate that both LPS and BDSF QS signal can elicit inflammatory responses in macrophages, whereas AHLs exhibit marked lower immunostimulatory activity.

**Fig. 1.**
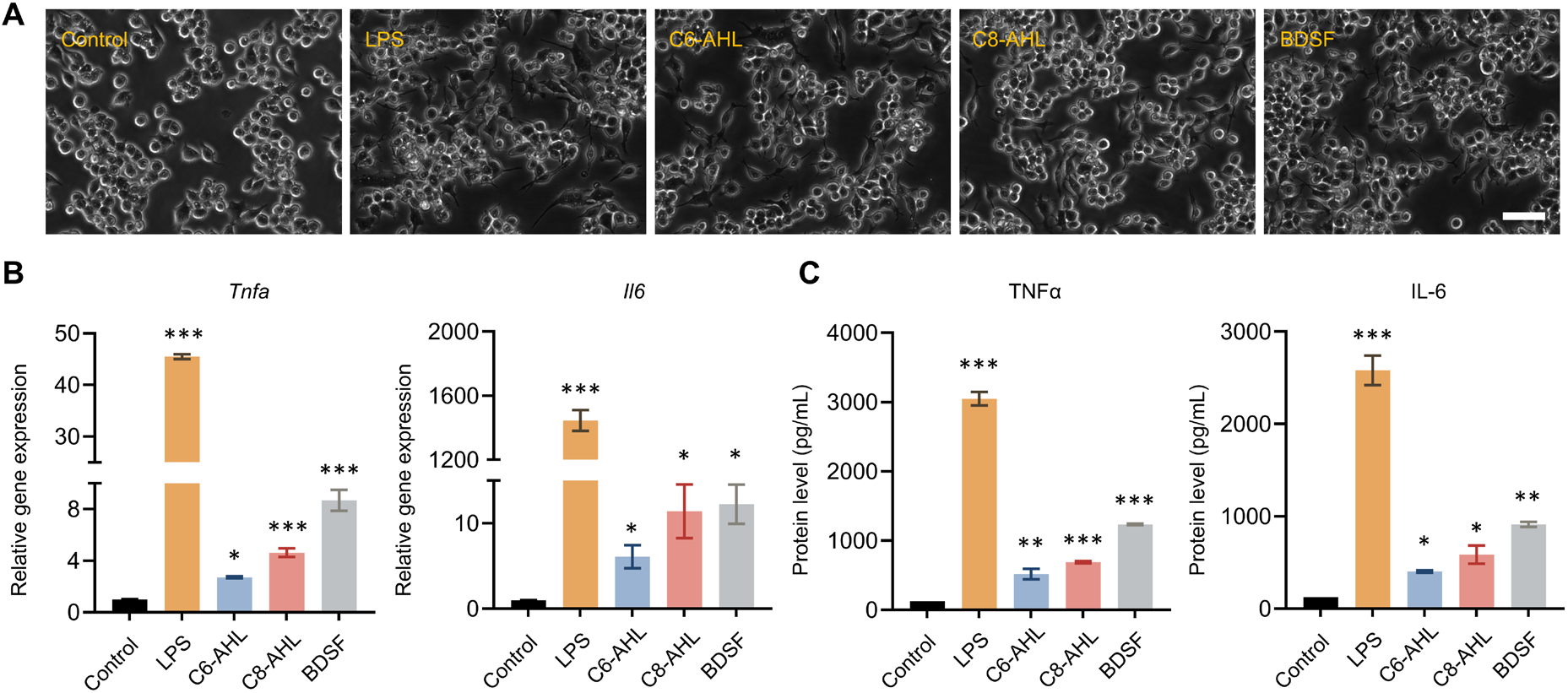
Quorum sensing molecules exert immunostimulatory effects on RAW264.7 cells. (**A**) The morphology of RAW264.7 cells treated with culture medium (Control), LPS, C6-AHL, C8-AHL, and BDSF for 24 hours was monitored. Scale bar, 50 μm. (**B**) The relative gene expression levels of *Tnfa* and *Il6* in RAW264.7 cells treated with culture medium, LPS, C6-AHL, C8-AHL, and BDSF, assayed by RT-qPCR. (**C**) The protein expression levels of TNFα and IL-6 in RAW264.7 cells treated with culture medium, LPS, C6-AHL, C8-AHL, and BDSF, assayed by ELISA. Data were presented as mean ± SEM. * *p* < 0.05, ** *p* < 0.01, and *** *p* < 0.001, compared with control group. Data were representative of at least three independent experiments with similar results.

### The immunostimulatory effects of BDSF in macrophages are independent of the canonical TLR4 signaling

To determine whether BDSF activates macrophages through the canonical Toll-like receptor 4 (TLR4) signaling pathway, we first performed microscale thermophoresis (MST) assays to assess potential interactions between BDSF and the extracellular domain (leucine-rich repeat, LRR) and intracellular domain (cytoplasmic Toll/interleukin-1 receptor, TIR) of TLR4. As shown in fig. S2, A and B, unlike LPS, BDSF did not induce measurable or concentration-dependent conformational changes in either TLR4 domain, indicating a lack of direct binding of BDSF to TLR4. To further verify the dispensable role of TLR4 in BDSF-induced inflammatory responses, we generated TLR4 knockout (KO) RAW264.7 cells and compared them to vector control

cells following BDSF treatment across a range of concentrations (fig. S2, C and D). BDSF elicited comparable upregulation of *Tnfa* and *Il6* transcripts in both vector and TLR4 KO cells (fig. S2E). Similarly, secretion of TNFα and IL-6 remained robust and indistinguishable in both vector and TLR4 KO cells upon BDSF stimulation (fig. S2F). Collectively, these results demonstrate that BDSF-mediated immune activation in macrophages is independent of TLR4 signaling, suggesting that alternative pattern recognition receptors (PRRs) or non-canonical sensing mechanisms mediate host recognition of BDSF.

### BDSF is a signal ligand of host EPAS1

To identify potential host sensors of BDSF, we performed limited proteolysis-small molecule mapping (LiP-SMap) using RAW264.7 macrophages treated with or without BDSF (table. S1). Differential protein profiling revealed 6 significantly enriched targets upon BDSF treatment, including PSMB5, TCP1, EPAS1, DNAJC2, NT5C2, and EIF2S1, with log_2_ fold changes (FC) of 8.66, 7.16, 6.75, -6.53, 6.50, and 6.27, respectively (Fig. 2, A and B). Each candidate protein was purified by affinity chromatography and tested for the direct interaction with BDSF using MST. No substantial binding was observed between BDSF and PSMB5, TCP1, DNAJC2, NT5C2, or EIF2S1 (fig. S3, A and B). For EPAS1, the protein domain architecture includes a basic helix-loop-helix (bHLH) conferring DNA binding and PAS domains conferring ligand-binding capacity (Fig. 2C). Interestingly, BDSF bound mouse recombinant EPAS1 with micromolar affinity, as demonstrated by both isothermal titration calorimetry (ITC, *K*_D_ value of 4.3 µM) and MST (*K*_D_ value of 14.5 µM) (Fig. 2, D to F). To assess the specificity of this interaction, we evaluated the binding affinity of EPAS1 to the AHLs QS, saturated BDSF isomer, or BDSF analog (DSF, *cis*-11-methyl-2-dodecenoic acid), using MST. None of these molecules showed binding capacity to EPAS1, indicating that BDSF is a specific ligand of host EPAS1 (fig. S4, A and B). Based on the previous report suggesting AHR as a potential host sensor for certain QS molecules (*23*), we also purified the binding domain of AHR and tested its interaction with BDSF, C6-AHL, and C8-AHL (fig. S4C). MST analysis showed that only C6-AHL and C8-AHL, but not BDSF, exhibited weak binding to AHR (fig. S4D). To examine whether the interaction between BDSF and EPAS1 is evolutionarily conserved, we first compared EPAS1 amino acid sequences across multiple species and found that the bHLH and PAS domains are highly conserved, particularly in *Mus musculus* and *Homo sapiens* (fig. S5, A and B). We then purified recombinant human EPAS1 protein and assessed its binding to BDSF. Both MST and ITC confirmed a direct interaction, with MST yielding a *K*_*D*_ value of 9.8 µM and ITC yielding a *K*_*D*_value of 5.3 µM (fig. S5, C and D).

**Fig. 2.**
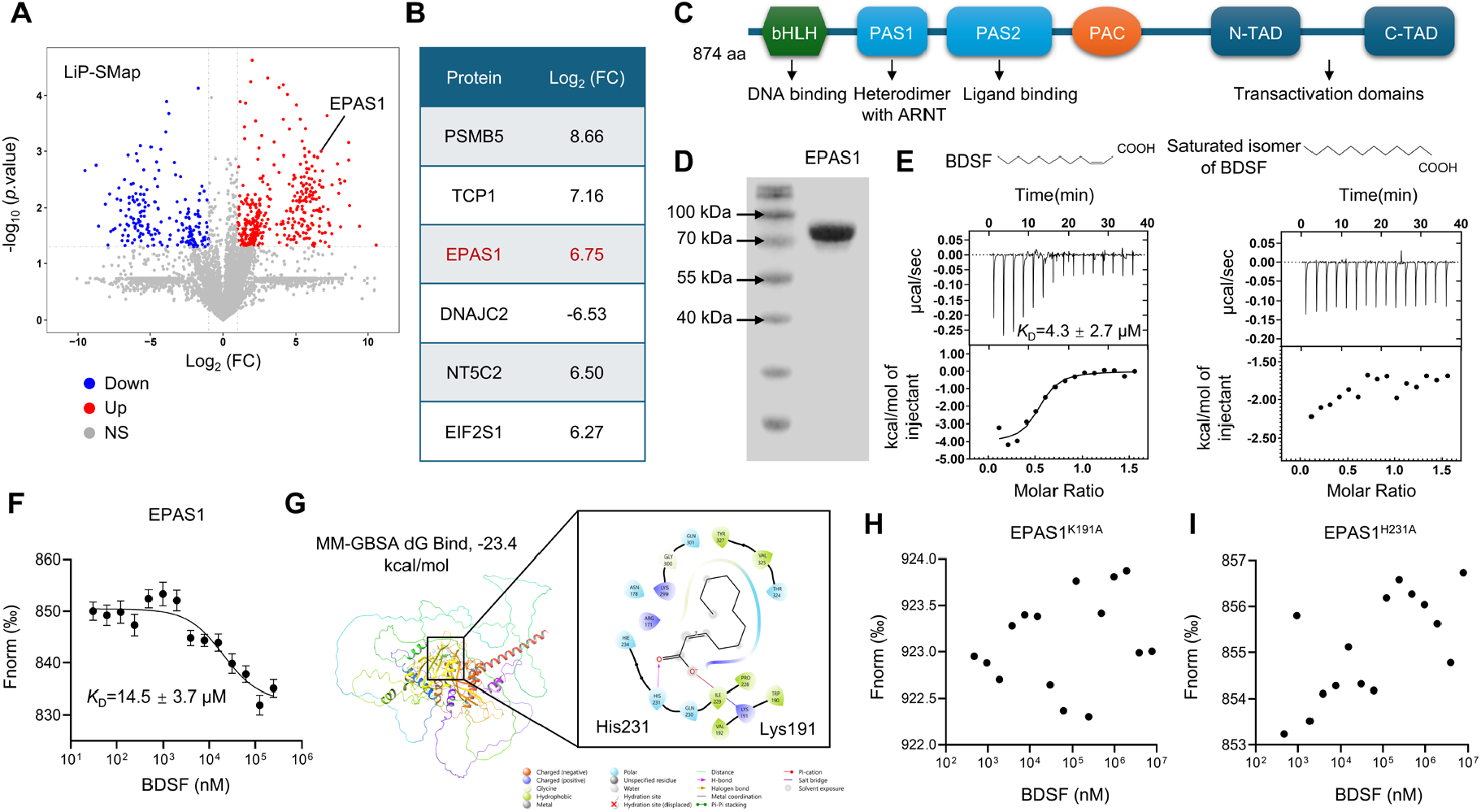
Identification of EPAS1 as a host receptor of BDSF. (**A**) Volcano plot illustrating differentially enriched proteins in RAW264.7 macrophages treated with BDSF compared to untreated controls, as identified by LiP-SMap analysis. (**B**) Top six BDSF-enriched host protein candidates identified by LiP-SMap. (**C**) The protein domain architecture of mouse EPAS1 includes a basic helix-loop-helix (bHLH) domain, Per-ARNT-Sim domains (PAS1 and PAS2), a PAS-associated C-terminal (PAC) motif, and N-terminal and C-terminal transactivation domains (N-TAD and C-TAD). (**D**) The SDS-PAGE analysis of purified EPAS1 protein, showing the purity and molecular weight of the recombinant protein. (**E**) Isothermal titration calorimetry (ITC) analysis of BDSF (left) or a saturated isomer of BDSF (right) binding to EPAS1. (**F**) MST analysis of BDSF binding to EPAS1. (**G**) Three-dimensional (3D) structural model of EPAS1 and chemical structure of BDSF. (**H-I**) MST analysis of BDSF binding to EPAS1 mutants K191A (H) and H231A (I). “Fnorm (‰)” represents the normalized fluorescence signal used to assess binding dynamics in the MST assay. Data were presented as mean ± SEM. Data were representative of at least three independent experiments with similar results.

To elucidate the molecular binding characteristics, we employed AlphaFold3 to predict the three-dimensional (3D) structure of mouse EPAS1, followed by binding site identification using Schrödinger software. Structure-based docking analysis implicated His231 (H231) and Lys191 (K191) in EPAS1 as critical residues for BDSF recognition (Fig. 2G). BDSF was predicted to form a hydrogen bond with H231 and a salt bridge with K191. To validate these findings, single-site mutants EPAS1^H231A^and EPAS1^K191A^ of EPAS1 were generated. MST analysis showed that mutations at either residue markedly weakened BDSF binding to EPAS1 (Fig. 2, H and I), supporting the essential role of H231 and K191 in the interaction between BDSF and EPAS1. Given that HIF family consist of HIF-1α, HIF-2α, and HIF-3α (*24*), we subsequently investigated whether BDSF could also bind to other HIF isoforms in addition to EPAS1. Docking analysis showed that the binding energies of BDSF with HIF-1α and HIF-3α were much higher than with EPAS1, supporting preferential binding to EPAS1 over other HIF isoforms (fig. S6, A and B). To further characterize the dynamics of EPAS1-BDSF interaction, we integrated molecular dynamics (MD) simulations with circular dichroism (CD) spectroscopy. The results revealed that BDSF maintained conformational flexibility within the EPAS1 binding site, engaging multiple interface regions while inducing a modest but reproducible shift in EPAS1 secondary structure from α-helices toward β-sheet and β-turn elements (fig. S7, A to D). Persistent intermolecular contacts, particularly involving the critical H231 residue, underscored high-affinity molecular interactions between EPAS1 and BDSF (fig. S7E). Collectively, these findings establish EPAS1 as a specific host receptor for BDSF and define a direct molecular interface for host sensing of QS signals.

### BDSF activates the MAPK pathway in an EPAS1-dependent manner

To elucidate the molecular mechanisms underlying BDSF-induced macrophage activation, we performed transcriptomic profiling of RAW264.7 cells with or without CRISPR/Cas9-mediated KO of *Epas1* (fig. S8). As shown in Fig. 3A, in response to BDSF treatment, *Epas1* deletion resulted in extensive transcriptional reprogramming and differential gene expression, compared to vector control cells (table. S2). Hierarchical clustering revealed distinct gene expression signatures between vector and EPAS1 KO groups (Fig. 3B). Kyoto encyclopedia of genes and genomes (KEGG) pathway enrichment analysis identified significant changes in several inflammation-related pathways in the absence of EPAS1, most notably the MAPK signaling pathway and TNF signaling pathways (Fig. 3C). Consistent with these transcriptomic changes, immunoblot analysis demonstrated that BDSF treatment induced the phosphorylation of p38, ERK1/2, and JNK, which were markedly diminished in EPAS1 KO macrophages (Fig. 3D). Furthermore, while BDSF induced a robust dose-dependent cytokine response in vector control cells, BDSF-induced secretion of TNFα and IL-6 was abolished in EPAS1 KO cells (Fig. 3E). To rule out confounding effects from cytotoxicity or altered EPAS1 expression, we confirmed that BDSF treatment did not affect cell viability up to 100 μM (fig. S9A) and had no impact on the mRNA or protein levels of EPAS1 across all concentrations of BDSF examined (fig. S9, B and C). Nevertheless, compared to the control group, BDSF treatment could promote the nuclear translocation of EPAS1 (fig. S9D), which further suggests the critical role of BDSF in regulating the transcription of EPAS1-targeted genes. Collectively, these results demonstrate that EPAS1 is required for BDSF-induced MAPK activation and expression of inflammatory cytokines in macrophages.

**Fig. 3.**
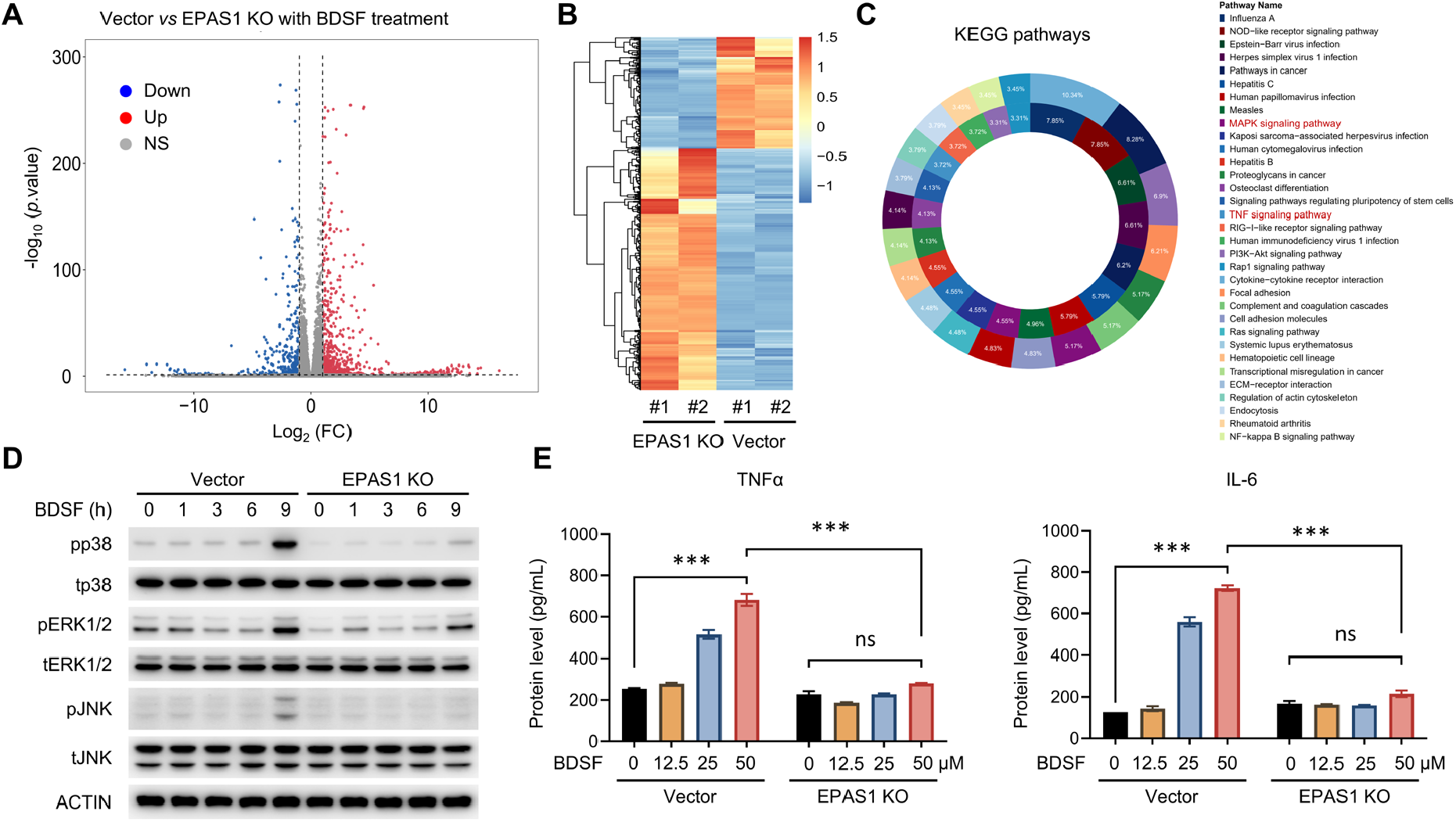
EPAS1-dependent cellular responses to BDSF. (**A**) Vector control and EPAS1-knockout (KO) RAW264.7 macrophages treated with BDSF were subjected to bulk RNA-seq analysis. The volcano plot displays differentially expressed genes (DEGs) between vector and EPAS1 KO cells under BDSF treatment. (**B**) Hierarchical clustering analysis of differentially expressed genes visualized as a heatmap. (**C**) KEGG pathway enrichment analysis of DEGs from RNA-seq, visualized as a doughnut chart. (**D**) Western blot analysis of MAPK pathway-related protein expression in vector and EPAS1 KO RAW264.7 cells following BDSF treatment for 0, 1, 3, 6, and 9 hours. (**E**) The protein expression levels of TNFα and IL-6 in RAW264.7 vector and EPAS1 KO cells treated with BDSF. Data were presented as mean ± SEM. * *p* < 0.05, ** *p* < 0.01, and *** *p* < 0.001, compared with the control or vector group as indicated. Data were representative of at least three independent experiments with similar results.

### EPAS1 controls the expression of target genes through direct promoter binding

As EPAS1 contains a bHLH domain predicted to be critical for DNA binding, we employed chromatin immunoprecipitation sequencing (ChIP-seq) to investigate the transcriptional mechanisms by which EPAS1 regulates macrophage inflammatory responses (table. S3). The proinflammatory cytokine genes *Tnfa* and *Il6* were among the identified targets (table. S3). Consistently, RT-qPCR confirmed the reduced *Tnfa* and *Il6* expression in EPAS1 KO cells compared to vector controls (Fig. 4A). To further elucidate the mechanism of EPAS1-mediated regulation, we first performed motif analysis (MEME-ChIP) on the ChIP-seq data, and identified a consensus EPAS1-binding site, 5’-CGCACGTAC-3’, within the promoters of these genes (Fig. 4B). To investigate the structural basis of EPAS1 interaction with this DNA binding motif, we conducted molecular docking and interaction mapping analyses (fig. S10). The docking data demonstrated that EPAS1 formed a stable complex with the promoter DNA motif (fig. S10A). Close-up visualization indicated that the residues Arg26 and Lys29 within the bHLH domain of EPAS1 serve as key contributors to DNA binding (fig. S10B). Upon addition of BDSF, the interaction profile changed notably, and several additional DNA-contacting residues, such as Arg27, Lys53, and His50, established a stronger interaction between EPAS1 and the DNA region in the promoter (fig. S10, C and D). Collectively, these results suggest that BDSF binding induces conformational rearrangements in EPAS1, promoting and diversifying its interaction with promoter DNA and potentially enhancing its transcriptional regulatory activity.

**Fig. 4.**
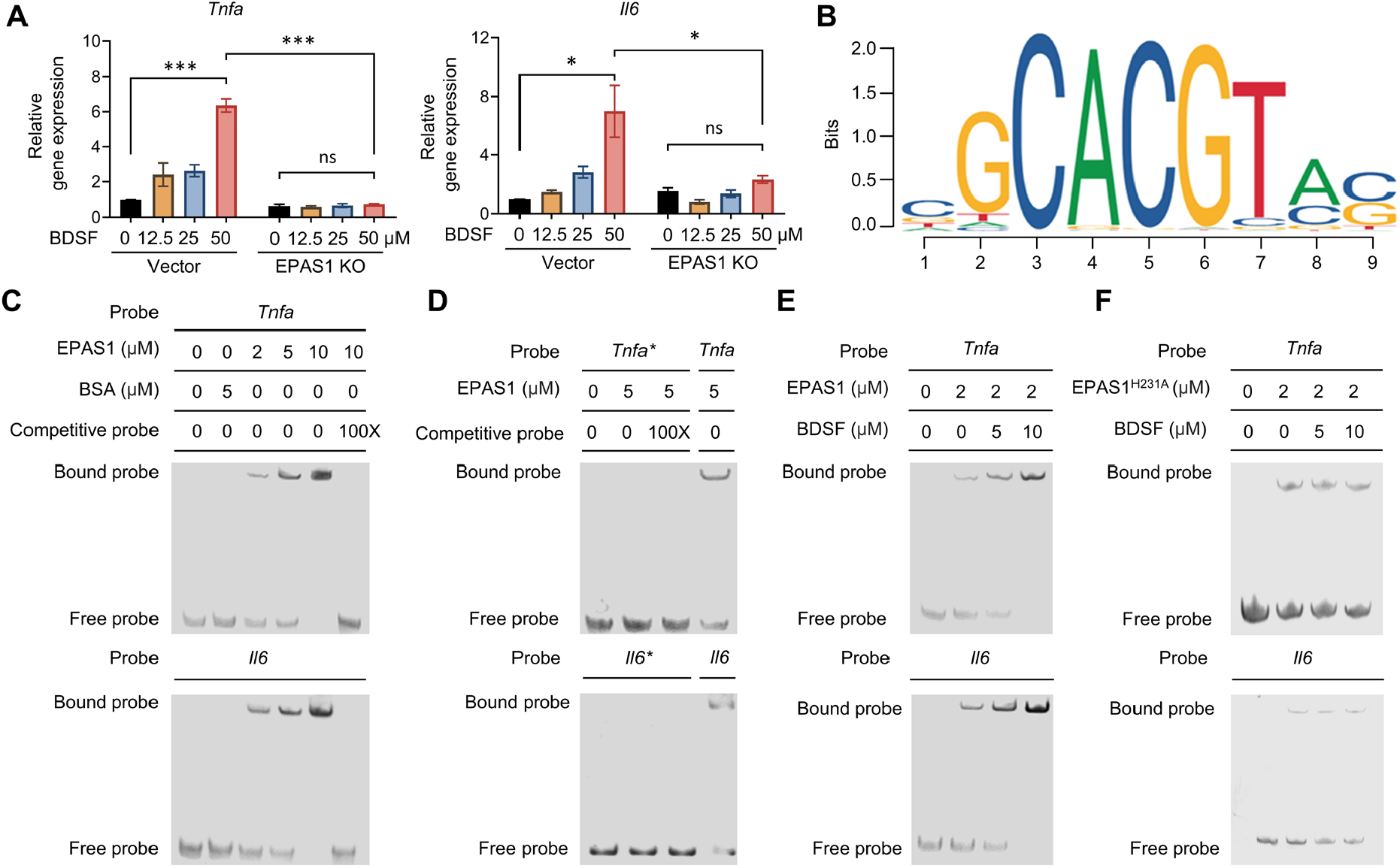
Regulation of target gene expression by EPAS1. (**A**) Relative mRNA levels of *Tnfa* and *Il6* in vector and EPAS1 KO RAW264.7 cells treated with BDSF. (**B**) Predicted EPAS1-binding sites in the promoter regions of target genes identified by ChIP-seq analysis. (**C-D**) Electrophoretic mobility shift assay (EMSA) demonstrating EPAS1 binding to the promoter regions of *Tnfa* and *Il6 in vitro* (C), an asterisk (*) indicates a mutated promoter fragment (D). (**E**) EMSA analysis of EPAS1 binding to *Tnfa* and *Il6* promoters in the presence of BDSF. (**F**) EMSA assay of EPAS1^H231A^ binding to *Tnfa* and *Il6* promoters in the presence of BDSF. Data were presented as mean ± SEM. * *p* < 0.05, ** *p* < 0.01, and *** *p* < 0.001, compared with the control or vector group as indicated. Data were representative of at least three independent experiments with similar results.

To validate the ChIP-seq findings, electrophoretic mobility shift assays (EMSAs) were performed. The results confirmed the direct binding of EPAS1 to the identified DNA motifs within the *Tnfa* and *Il6* promoters (Fig. 4C). Increasing concentrations of EPAS1 resulted in progressively increasing amounts of DNA-protein complexes, whereas competition with excess unlabeled probe abolished the binding. Furthermore, deletion of the binding site from the *Tnfa* and *Il6* promoter regions abrogated complex formation (Fig. 4D), demonstrating that this specific sequence is essential for EPAS1 binding. Given that transcription of *Tnfa* and *Il6* is closely regulated by MAPK signaling pathways, we further analyzed the ChIP-seq dataset to determine whether EPAS1 also directly targets upstream regulators of the MAPK cascade. Gene ontology (GO) enrichment analysis revealed a significant overrepresentation of MAPK-related processes among EPAS1-associated gene promoters (fig. S11A). Among the candidate genes, *Map2k2, Map3k7*, and *Map3k8*, which encode core MAPK regulators, were selected for further validation (fig. S11B). MST assays demonstrated high-affinity interactions between EPAS1 and the promoter regions of these genes (fig. S11C). Consistent with these results, EMSA results showed a concentration-dependent formation of EPAS1-DNA complexes that were effectively competed by excess unlabeled probe (fig. S11D). Importantly, deletion of the EPAS1 binding motif within the *Map2k2* promoter resulted in a loss of complex formation (fig. S11E). Taken together, these findings demonstrate that EPAS1 not only directly binds to the promoters of proinflammatory cytokine genes, but also to those of key MAPK pathway regulators, thereby regulating inflammatory signaling at multiple levels.

### BDSF enhances the binding of EPAS1 to target gene promoters

To assess whether BDSF modulates the transcriptional activity of EPAS1, we conducted EMSA assays in the presence or absence of BDSF. BDSF enhanced EPAS1 binding to the *Tnfa* and *Il6* promoters in a concentration-dependent manner (Fig. 4E). Similarly, BDSF increased the binding affinity between EPAS1 and *Map2k2* gene promoter, and promoted the transcription of *Map2k2, Map3k7*, and *Map3k8* in an EPAS1-dependent manner (fig. S11, F and G). Notably, mutation of BDSF-binding site (EPAS1^H231A^) eliminated the effect of BDSF on the ability of EPAS1 to bind the target gene promoters (Fig. 4F). To further investigate the effect of EPAS1 inhibition on BDSF-induced transcriptional responses, we performed molecular docking between EPAS1 and its inhibitor PT2399 (*9*) (fig. S12A). The docking analysis showed that EPAS1 inhibitor bound to EPAS1 with high affinity (-59.0 kcal/mol), indicating a strong inhibitory interaction (fig. S12A). Supporting this prediction, pharmacological inhibition of EPAS1 with PT2399 significantly reduced BDSF-induced expression of the inflammatory genes *Tnfa* and *Il6* (fig. S12B). Similarly, the EPAS1 inhibitor nearly abolished the BDSF-induced upregulation of MAPK regulator genes *Map2k2, Map3k7*, and *Map3k8* (fig. S12C). Previous studies have demonstrated that EPAS1 forms the heterodimer with ARNT to initiate gene expression (*4*). To further examine whether BDSF alters the structural conformation of the EPAS1-ARNT complex, we performed CD spectroscopy. As shown in fig. S13, the CD spectra of EPAS1-ARNT displayed features consistent with β-sheet and random coil-rich conformations. In contrast, addition of BDSF caused a marked spectral shift, indicating that BDSF binding significantly increased α-helical content while reducing random coil proportion (fig. S13, B and C). Collectively, these results demonstrate that EPAS1 is essential for BDSF-driven activation of inflammatory and MAPK signaling genes, and its pharmacological blockade markedly dampens the transcriptional responses.

### BDSF promotes *B. cenocepacia* and *A. baumannii*-triggered inflammatory responses through EPAS1

To investigate whether BDSF can potentiate macrophage inflammatory responses to bacterial pathogens, we first assessed its effect in the context of *B. cenocepacia* strain H111 exposure. In RAW264.7 cells, co-stimulation with BDSF and strain H111 cells significantly enhanced *Tnfa* and *Il6* transcription compared to either stimulus alone (Fig. 5A), accompanied by increased phosphorylation of p38, ERK1/2, and JNK (Fig. 5B). These transcriptional and signaling enhancements were markedly reduced in EPAS1 KO cells, indicating that BDSF amplifies H111-induced inflammatory responses in an EPAS1-dependent manner. The immunoregulatory effect of BDSF was extended to *A. baumannii*, another clinically relevant Gram-negative pathogen without producing BDSF. Similar to the findings from H111 infections, combined stimulation with BDSF and *A. baumannii* markedly increased *Tnfa* and *Il6* expression and activation of MAPKs in the vector control cells, but not in EPAS1 KO macrophages (Fig. 5, C and D). Cytokine quantification further confirmed that elevated TNFα and IL-6 secretion were observed in response to combined stimulation of BDSF with either pathogen, and the enhanced response was abrogated again in the absence of EPAS1 (Fig. 5, E and F). Together, these findings demonstrate that BDSF amplifies macrophage inflammatory responses to bacteria regardless of whether they produce BDSF, via an EPAS1-dependent mechanism.

**Fig. 5.**
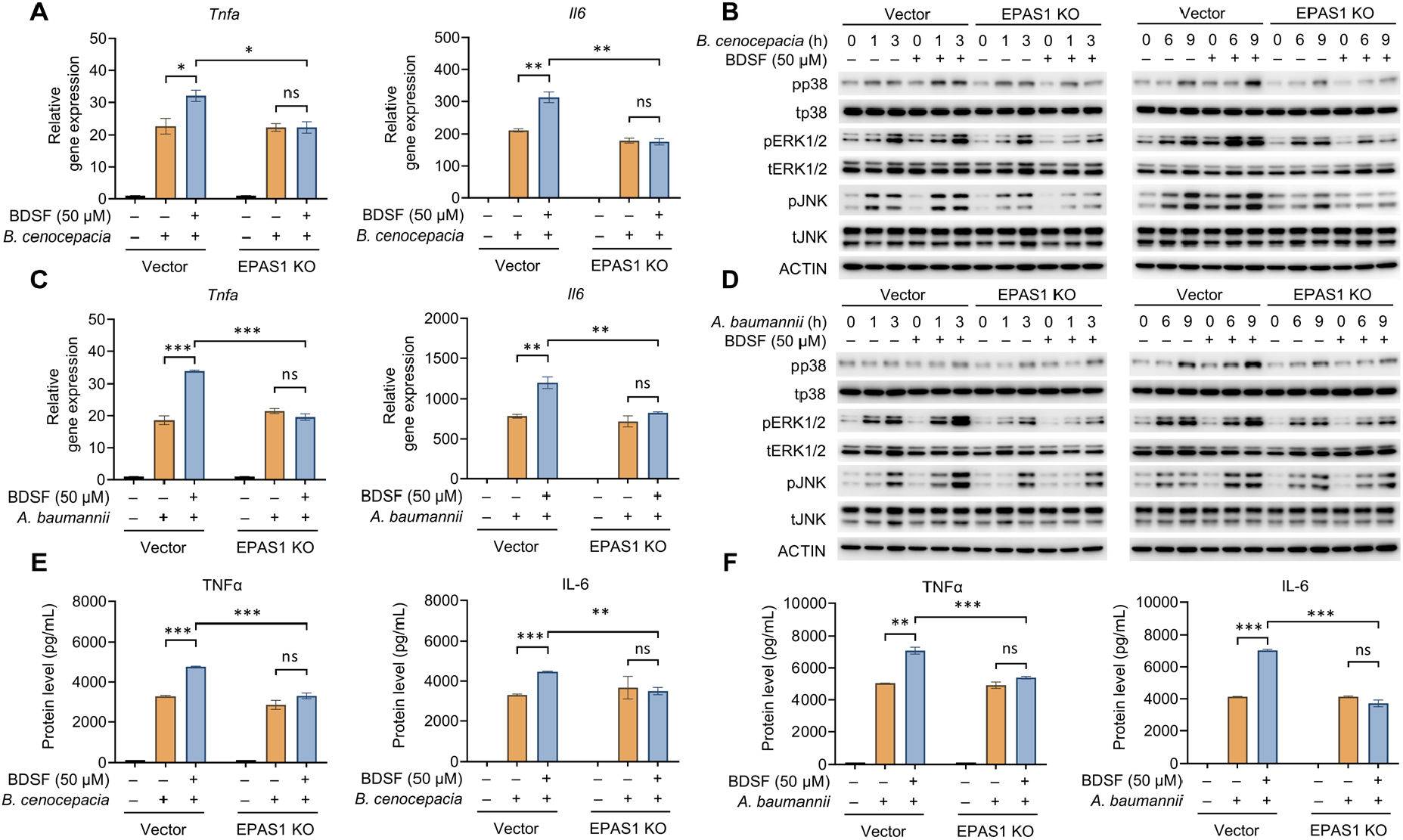
EPAS1-dependent effects of BDSF on inflammatory responses to *B. cenocepacia* and *A. baumannii* in RAW264.7 macrophages. (**A**) Relative gene expression levels of *Tnfa* and *Il6* in vector and EPAS1 KO RAW264.7 cells following *cenocepacia* stimulation, with or without BDSF pretreatment. (**B**) Western blot analysis of MAPK pathway-related protein expression in vector and EPAS1 KO RAW264.7 cells after *B. cenocepacia* stimulation, with or without BDSF pretreatment. (**C**) Relative gene expression levels of *Tnfa* and *Il6* in vector and EPAS1 KO RAW264.7 cells following *A. baumannii* stimulation. (**D**) Western blot analysis of MAPK pathway-related protein expression in vector and EPAS1 KO after *A. baumannii* stimulation. (**E**) Protein levels of TNFα and IL-6 in vector and EPAS1 KO RAW264.7 cells stimulated with *B. cenocepacia* and BDSF. (**F**) Protein levels of TNFα and IL-6 in vector and EPAS1 KO RAW264.7 cells stimulated with *A. baumannii* and BDSF. Data were presented as mean ± SEM. * *p* < 0.05, ** *p* < 0.01, and *** *p* < 0.001, compared with the control or vector group as indicated. Data were representative of at least three independent experiments with similar results.

### Inoculation of BDSF mitigates *B. cenocepacia* and *A. baumannii* infections *in vivo*

To determine whether the immunostimulatory functions of BDSF confer protection against bacterial infections *in vivo*, we established a murine pneumonia model via intratracheal injection of *B. cenocepacia* or *A. baumannii*, with prior BDSF treatment (Fig. 6A). Mice pretreated with BDSF alone for 4 days exhibited only mild inflammation by histology and cytokine measurements, with no significant changes in body weight or survival compared to controls (Fig. 6, B to F). Strikingly, BDSF pretreatment markedly improved survival in mice upon H111 infection (Fig. 6, B and C). While all untreated mice succumbed to infection by day 7, 3 out of 7 BDSF-pretreated mice (42.9% survival) survived until day 8 (Fig. 6C). Histological analysis revealed reduced inflammatory cell infiltration and tissue injury in the lung, accompanied by significantly lower expression of the inflammatory cytokines *Tnfa* and *Il6* in BDSF-pretreated mice compared to mice without treatment (Fig. 6, E and F). In addition, similar protective effects were observed in a separate cohort infected with *A. baumannii* (Fig. 6, B to F). These findings suggest that BDSF pretreatment enhances host immune readiness, attenuates hyperinflammatory responses, and confers protection against lethal bacterial challenge.

**Fig. 6.**
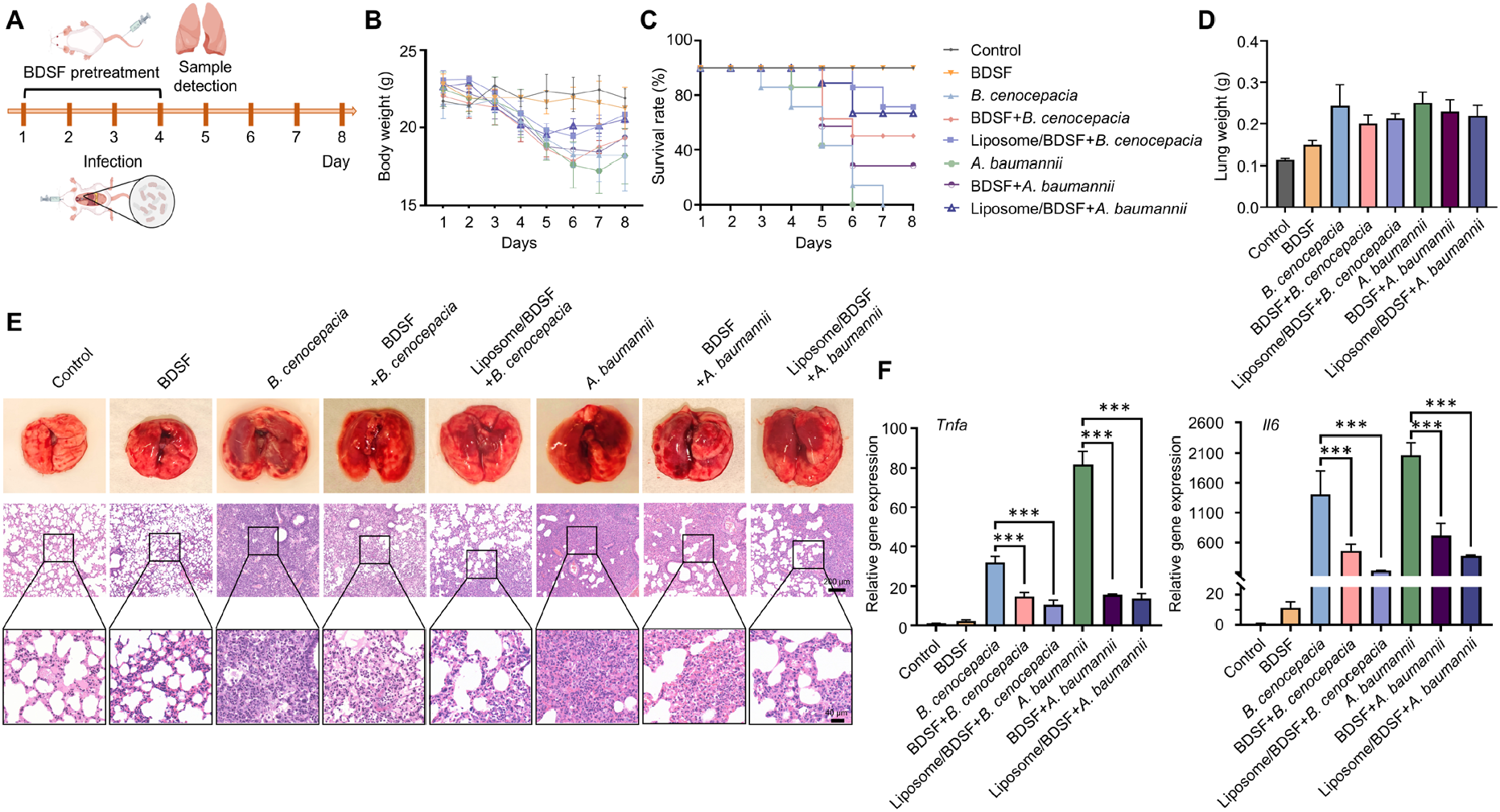
*In vivo* effects of BDSF on pulmonary inflammation in *B. cenocepacia*- and *A. baumannii*-infected mice. (**A**) Schematic diagram illustrating pulmonary infection with *B. cenocepacia* and *A. baumannii* bacteria in mice. (**B**) Body weight changes of mice upon bacterial infections. (**C**) Survival curves of mice post bacterial infections. (**D**) Lung weight of mice infected with *B. cenocepacia* and *A. baumannii*. (**E**) Representative microscopic images of lung tissues and histopathological examination. (**F**) Relative mRNA levels of *Tnfa* and *Il6* in lung tissues from mice with bacterial infections. Data were presented as mean ± SEM. * *p* < 0.05, ** *p* < 0.01, and *** *p* < 0.001. Data were representative of at least three independent experiments with similar results.

In recent years, lipid nanoparticles (LNPs) have become an important delivery platform due to their ability to enhance drug stability, safety, and delivery efficiency. To further evaluate the clinical potential of BDSF-based improvement of host immune responses, we encapsulated BDSF in liposomes (Liposome/BDSF) to improve its *in vivo* bioavailability and immunostimulatory effects without inducing cytotoxicity in immune cells (fig. S14). Pretreatment with Liposome/BDSF significantly improved survival in bacteria-infected mice, as evidenced by a survival rate of 71.4% in the Liposome/BDSF group infected with *B. cenocepacia* or *A. baumannii* (Fig. 6C). Histopathological analysis revealed substantially reduced inflammatory cell infiltration and tissue damage in the lungs of Liposome/BDSF-pretreated mice, together with decreased expression of the inflammatory cytokines *Tnfa* and *Il6* (Fig. 6, D to F). We hypothesize that this protective effect is mediated by moderate immune stimulation and EPAS1 activation, which enhances bacterial clearance *in vivo*. To test this, we performed bacterial load assays and flow cytometric analyses on the inflamed lung tissues. Mice pretreated with Liposome/BDSF and subsequently infected with either *B. cenocepacia* or *A. baumannii* exhibited significantly lower lung bacterial loads and a higher proportion of pulmonary macrophages among leukocytes, compared to untreated infected controls (fig. S15). These results suggest that BDSF-driven macrophage activation promotes bacterial clearance, thereby reducing tissue injury. In addition, to further assess the safety and tolerability of the LNPs-based delivery strategy, we conducted histopathological examination of kidney and colon tissues upon BDSF treatment. Neither BDSF nor Liposome/BDSF was associated with carcinogenicity or colonic enteritis at the tested dosage for 7 days *in vivo* (fig. S16). Consistently, the expression levels of proliferation marker *Mki67* in the kidney and mucin gene *Muc2* in the colon were comparable across treatment groups, further supporting the absence of adverse effects (fig. S16, B to D). Together, these findings demonstrate that BDSF induces EPAS1-mediated immune-protection against bacterial infection while maintaining a favorable safety profile, suggesting that BDSF and its derivatives could serve as promising immunomodulatory agents for preventing or treating bacterial infections.

## Discussion

The capacity of hosts to perceive and interpret microbial QS signals represents a promising research field to be explored (*23, 25*). Bacterial QS systems, as exemplified by *P. aeruginosa* and *B. cenocepacia*, orchestrate collective behaviors such as biofilm formation, virulence, and antibiotic resistance (*16, 19*). While QS is fundamental for bacterial adaptation, recent studies showed that hosts can also sense and respond to these microbial signals, although the underlying receptors and pathways have remained largely undefined. Notably, the AHR was previously identified as a highly conserved, ligand-dependent transcription factor that senses environmental toxins and endogenous ligands, including phenazines from *P. aeruginosa* and naphthoquinone phthiocol from *M. tuberculosis* (*23, 26*). Upon binding these ligands, AHR initiates transcriptional programs to regulate cytokine production, antibacterial defenses, and virulence factor degradation, establishing it as the first known intracellular pattern recognition receptor (PRR) that monitors bacterial quorum status to dynamically modulate immune responses (*23, 26*). In this study, we identify EPAS1 as a previously unrecognized sensor for the bacterial QS molecule BDSF, elucidating a new mechanism by which microbial signaling metabolites are detected by host immune cells. Our results indicate that BDSF-driven activation of EPAS1 enhances macrophage inflammatory responses and confers robust protection against both *B. cenocepacia* and *A. baumannii* infections *in vivo*, supporting a concept of gentle immunity whereby BDSF-EPAS1 axis strengthens host defense without provoking excessive inflammation. The evolutionary adaptation underlying the specificity of EPAS1 for BDSF from *B. cenocepacia* highlights a finely tuned molecular interplay between host transcriptional regulators and bacterial metabolites. In contrast to classical PRRs, which respond to invariant pathogen-associated molecular patterns (*27*), EPAS1 appears to function as a sensor of microbial behavior, enabling context-dependent responses that are temporally and functionally distinct from those triggered by structural components like LPS or peptidoglycan (*28, 29*). Taken together, this study positions EPAS1 as a novel intracellular sensor for bacterial QS signals, fundamentally broadening the conceptual framework of host-pathogen interkingdom communication and providing a foundation for metabolite-guided strategies in immunotherapy, immunomodulator development, and infectious disease management.

Mechanistically, we demonstrate that EPAS1 serves as a crucial transcriptional regulator linking recognition of microbial signals to the activation of host inflammatory pathways. EPAS1 directly associates with the promoter regions of MAPK pathway regulators, including *Map2k2, Map3k7*, and *Map3k8*, and enhances their transcriptional expression in response to BDSF sensing. Activation of the MAPK signaling cascade subsequently leads to the production of inflammatory cytokines such as TNFα and IL-6 (*30*). These cytokines, in turn, activate the MAPK signaling pathway, creating a positive feedback circuit that amplifies innate immune cell activation (*31, 32*). In addition, we confirmed the binding of EPAS1 to the promoters of *Tnfa* and *Il6*, suggesting the direct regulation of EPAS1 in the transcription of cytokines (Fig. 4C). This interconnected regulatory network provides a molecular basis for the observed enhancement of immune responses following BDSF exposure. Importantly, genetic ablation or pharmacological inhibition of EPAS1 abolishes both MAPK activation and cytokine production in response to BDSF or bacterial challenge, confirming the central role of EPAS1 in this signaling axis (Fig. 3 and 6). Together, these results establish EPAS1 as an upstream regulator of MAPK signaling and infection-induced innate immunity. The immunomodulatory function of BDSF is further substantiated by its ability to augment macrophage responses to both *B. cenocepacia* and *A. baumannii*, two clinically relevant and multidrug-resistant pathogens (*33, 34*). Unlike robust immunostimulants, such as LPS, which can precipitate overwhelming and potentially damaging inflammatory responses (*35*), BDSF triggers a moderate upregulation of key inflammatory cytokines, including TNFα and IL-6. This intermediate activation state is sufficient to prime macrophages for enhanced pathogen recognition and clearance (Fig. 6). By harnessing BDSF or related agonists as prophylactic agents, it may be possible to selectively activate innate immune pathways, providing broad-spectrum resistance to bacterial pathogens while minimizing the risk of cytokine storm. This work thus provides a compelling blueprint for the rational design of immunomodulatory interventions rooted in the principles of gentle immune activation.

EPAS1 is a pleiotropic transcription factor centrally involved in a variety of physiological and pathological processes, including hypoxia adaptation, oxidative stress regulation, cell proliferation, apoptosis, and angiogenesis (*10, 36, 37*). The activity of EPAS1 is critical in developmental and homeostatic settings; However, aberrant or excessive EPAS1 activation has been extensively implicated in cancers. For instance, EPAS1 drives angiogenesis via vascular endothelial growth factor (VEGF) upregulation and has been implicated in both cardiovascular pathology and tumor progression (*38*). Notably, pharmacological inhibition of EPAS1 with small molecules such as PT2399 can suppress tumor progression in preclinical models of kidney carcinoma (*9*). In the present study, we uncover a novel immunological function for EPAS1 as an intracellular sensor for bacterial QS molecules, which enables EPAS1 to modulate innate immunity by enhancing the expression of inflammatory cytokines and promoting antibacterial defense. This previously unappreciated EPAS1-mediated pathway highlights a new axis of host-pathogen communication, positioning EPAS1 as a promising target for host-directed therapy against multidrug-resistant bacterial infections. Notably, BDSF-mediated preconditioning conferred significant protection in mouse models of pneumonia caused by *B. cenocepacia* and *A. baumannii*, with no observable tissue damage or carcinogenicity in the kidney and colon after short-term exposure. Our findings not only expand the known functional landscape of EPAS1 but also underscore the necessity for rigorous safety assessment prior to clinical translation of EPAS1-targeted interventions, especially in light of its broad physiological relevance. However, given the multifaceted roles of EPAS1 in essential cellular functions and its established involvement in tumorigenesis, there is a clear need for comprehensive long-term biosafety studies to fully evaluate the safety and therapeutic window of this approach.

In summary, this study establishes EPAS1 as a selective, evolutionarily conserved, and functionally significant host receptor for the bacterial QS molecule BDSF, thereby unveiling a direct molecular link between microbial communication signals and host immune response. Our work demonstrates that BDSF binding to the PAS domain of EPAS1 triggers a cascade of transcriptional events culminating in the activation of MAPK signaling and protective immune responses. By defining this unique mode of host-pathogen interaction, we expand the repertoire of immune sensors and reveal a previously unknown mechanism by which microbial QS can reprogram host transcriptional networks. This new understanding uncovers an additional layer of host-microbe interaction and sets the stage for the development of bacterial QS-guided strategies in immunotherapy and infectious disease management.

## Materials and Methods

### Bacterial culture

*B. cenocepacia* H111 was cultured in lysogeny broth (LB, 5 g/L yeast extract (Biosharp, Beijing, China), 10 g/L tryptone (Biosharp), and 10 g/L NaCl) at 37 °C. For solid media, 15 g/L agar (Biosharp) was added. *A. baumannii* was grown in the LB broth and agar plates at 37 °C.

### Preparation of the extract of *B. cenocepacia* culture supernatant (BCS)

*B. cenocepacia* H111 was streaked onto LB agar plates for isolation. A single colony was selected and inoculated into 10 mL of LB broth, incubated at 37 °C, and shaken overnight at 220 rpm. The overnight culture was then used to inoculate 1 L of LB broth and cultured at 37 °C with shaking at 200 rpm until an OD_600_ of 3.0 was reached. The bacterial culture was centrifuged at 12,000 rpm for 5 minutes at 4 °C, and the supernatant was collected. Hydrochloric acid was added to adjust the pH to 4, followed by the addition of an equal volume of ethyl acetate. The mixture was vigorously shaken, transferred to a separating funnel for phase separation, and the upper organic phase was collected. The ethyl acetate was evaporated using a rotary evaporator (RE-52AA) in a fume hood, and the dried extract was dissolved in chromatography-grade methanol, filtered through a 0.22 μm filters, and stored at -20 °C.

### Statistical analysis

Data were presented as mean ± standard error of the mean (SEM). No data were excluded from the analysis. GraphPad Prism (version 8.3.0.538) was used for statistical analysis and *p* < 0.05 was considered statistically significant. Comparisons between two groups were conducted by two-tailed and nonparametric Student’s *t*-tests. For comparison of 3 or more groups, one-way ANOVA followed by Turkey’s multiple comparison test was used. * *p* < 0.05, ** *p* < 0.01, *** *p* < 0.001, and ‘ns’ indicated no significant difference. All experiments presented were repeated at least 3 times with similar results. Additional Materials and Methods are available in Supplementary Information.

### Reporting summary

Further information on research design is available in the Nature Portfolio Reporting Summary linked to this article.

## Data availability

The main data supporting the results of this study are available within the paper and Supplementary Information. All the raw and analyzed data generated during the study are available from the corresponding authors on reasonable request.

## Author contributions

HL, XK, BC, YZ, and YD conceived the study and wrote the manuscript. HL, XK, BC, XL, XL, MW, and LEW performed the experiments. HL, XK, BC, XL, XL, MW, LEW, XC, WQ, LHZ, YZ, and YD analyzed and interpreted the data. YZ and YD revised the manuscript, obtained the funding, and supervised the whole study. All authors reviewed and edited the manuscript.

## Supplementary Materials

Materials and Methods

Figs. S1 to S16

Tables S1 to S5

References

