## Supplemental Material for "EPAS1 regulates host antibacterial defense in response to bacterial quorum sensing"

**Materials and Methods**

**Cell culture and treatment**

RAW264.7, a mouse macrophage cell line, was obtained from the American Type Culture Collection (ATCC, Manassas, VA, USA) and cultured in Dulbecco's Modified Eagle Medium (DMEM) supplemented with 10% (v/v) fetal bovine serum (FBS), 100 U/mL penicillin, and 100 μg/mL streptomycin. RAW264.7 cells were authenticated by STR profiling and tested mycoplasma-negative. The cells were maintained in a humidified incubator (ESCO) at 37 °C with 5% $CO_2$ and sub-cultured biweekly. At 50% confluence in six-well plates, RAW264.7 cells were treated with 50 μM C6-AHL, C8-AHL, or BDSF, while lipopolysaccharide (LPS) at 1 μg/mL served as a positive control. To assess the cellular response to bacterial infections, RAW264.7 cells were infected with *B. cenocepacia* H111 or *A. baumannii* at $10^5$ CFU/mL, either in the absence or presence of 50 μM BDSF. Following infection and treatment, cell morphology was monitored under Olympus microscope, and the cells were harvested for gene and protein expression analyses. To evaluate the impact of EPAS1 inhibition on inflammatory responses, RAW264.7 cells were treated with the selective EPAS1 inhibitor PT2399 (Aladdin Scientific, Riverside, CA, USA).

**Limited proteolysis small molecule mapping (LiP-SMap) analysis**

LiP-SMap assay was performed to identify protein regions conformationally altered upon ligand binding. Briefly, RAW264.7 cell lysates were prepared in a native buffer and clarified by centrifugation. Equal protein amounts were incubated with BDSF or control for 30 minutes at room temperature. Limited proteolysis was initiated by adding nonspecific protease and allowed to proceed for 5 minutes at room temperature to achieve partial digestion; reactions were quenched with protease inhibitor cocktail and rapid heat denaturation. Subsequently, peptides were desalted and analyzed by liquid chromatography-tandem mass spectrometry (LC-MS/MS). Raw data were processed against the species proteome database with a target-decoy strategy. Peptide intensities were normalized across runs and differential peptide abundance between ligand and vehicle conditions was tested using linear models.

**Bulk RNA-seq analysis**

Total RNA was isolated from BDSF-treated RAW264.7 vector control and EPAS1 KO cells using TRIzol reagent (Invitrogen), followed by poly(A)+ mRNA enrichment. Bulk RNA-seq libraries were prepared with the Illumina RNA Sequencing Preparation Kit and sequenced on the Illumina HiSeq platform. Reads were aligned to the *Mus musculus* reference genome (GRCm39/mm39) with HISAT2 using default parameters. Transcript assembly and quantification were performed with StringTie (v2.1.4), and fragments per kilobase of transcript per million mapped reads (FPKM) values were calculated to quantify gene-level mRNA expression. Differential expression analysis was conducted using the DESeq2 package in R software to identify differentially expressed genes (DEGs). All analyses and visualizations, including Kyoto Encyclopedia of Genes and Genomes (KEGG) pathway enrichment, heatmaps, and volcano plots, were performed in R (v4.3.2) within RStudio.

**Generation of RAW264.7 KO cells**

To generate EPAS1 or TLR4 KO cells, single guide RNAs (sgRNAs) were designed using the MIT online CRISPR design tool and synthesized by Genscript (Nanjing, China). The sgRNAs were complexed with Cas9 protein *in vitro* to form RNP complexes, and the resulting complexes were introduced into RAW264.7 cells by electroporation. Single-cell-derived colonies were isolated and screened for gene disruption by Sanger sequencing and validated at the protein level by Western blotting.

**Protein expression and purification**

Using mouse (*Mus musculus*) cDNA or genome from RAW264.7 cells and human (*Homo sapiens*) cDNA from A549 cells (ATCC) as a template, the open reading frame of the target gene was amplified in two segments. The PCR products were gel-purified and subsequently ligated into the linearized pET-28a (+) plasmid. Detailed information about plasmids were listed in table. S4. The fusion plasmid was then transformed into *E. coli* BL21 (DE3) and cultured in LB broth supplemented with kanamycin at 37 °C. When the culture broth reached an $OD_{600}$ of 0.6, isopropyl-β-D-thiogalactoside (IPTG, Sigma-Aldrich, Missouri, USA) was added to a final concentration of 0.5 mM, and the culture was incubated at 16 °C for 10 hours.

Following protein expression, bacterial cells were harvested via centrifugation at 4,000 rpm for 15 minutes at 4 °C and resuspended in phosphate-buffered saline (PBS). The cell suspension was subjected to ice-cooled ultrasonication for 20 minutes using an ultrasonic homogenizer (SCIENTZ, JY92-IIN) to achieve lysis. Cellular debris was removed by centrifugation at 8,000 rpm for 30 minutes at 4 °C, and the resulting supernatant was loaded onto a Ni-NTA affinity chromatography column. Non-specifically bound proteins were eliminated by washing the column with a buffer containing 50 mM sodium phosphate, 0.3 M NaCl, and 20 mM imidazole (pH 8.0). The target His-tagged protein was subsequently eluted using an elution buffer composed of 50 mM sodium phosphate, 0.3 M NaCl, and 250 mM imidazole (pH 8.0). The eluted protein fraction was concentrated using an ultrafiltration tube (Millipore, Billerica, MA, USA), and purity was assessed by SDS-PAGE analysis.

**Chromatin immunoprecipitation sequencing (ChIP-seq) assay**

Chromatin immunoprecipitation (ChIP) assays were performed as previously described by Wuhan IGENEBOOK Biotechnology (*1*). Briefly, RAW264.7 cells ($10^6$) were washed twice with cold PBS and cross-linked with 1% formaldehyde for 10 minutes at room temperature. Cross-linking was quenched by the addition of 125 mM glycine. Cells were then lysed in a buffer containing 50 mM Tris-HCl (pH 8.0), 10 mM EDTA, 1% SDS, and a protease inhibitor cocktail. Chromatin was isolated on ice and subsequently sheared by sonication to an average DNA fragment size of 200-500 bp. A 20 µL aliquot of chromatin was reserved at -20 °C as input control, and 100 µL was subjected to immunoprecipitation using 10 µg of anti-His antibody (ab9108, Abcam, Cambridge, UK) at 4 °C overnight. The following day, 30 µL of protein G magnetic beads (10004D, Thermo Fisher Scientific) were added, and samples were incubated for an additional 3 hours. Beads were sequentially washed: once with low-salt buffer (20 mM Tris-HCl, pH 8.1; 50 mM NaCl; 2 mM EDTA; 1% Triton X-100; 0.1% SDS); twice with high-salt buffer (10 mM Tris-HCl, pH 8.1; 250 mM LiCl; 1 mM EDTA; 1% NP-40; 1% sodium deoxycholate); and twice with TE buffer (10 mM Tris-HCl, pH 7.5; 1 mM EDTA). Bound DNA was eluted in 300 µL of elution buffer (100 mM $NaHCO_3$, 1% SDS), followed by RNase A treatment (8 µg/mL, 6 hours at 65 °C) and proteinase K digestion (345 µg/mL, overnight at 45 °C). The resulting immunoprecipitated DNA was used to construct ChIP-seq libraries using the NEXTFLEX® ChIP-Seq Library Prep Kit for Illumina® (NOVA-5143-02, Bioo Scientific) and sequenced on an Illumina NovaSeq 6000 platform (paired-end 150 bp).

Raw reads were quality-filtered using Trimmomatic (v0.36), and clean reads were aligned to the RAW264.7 reference genome using BWA (v0.7.15). PCR duplicates were removed with Samtools (v1.3.1). Peak calling was conducted using MACS2 (v2.1.1.20160309) with default parameters (bandwidth: 300 bp; model fold: 5-50; *q*-value cutoff: 0.05). Peaks were assigned to genes based on proximity to the transcription start site (TSS). Motif enrichment analysis within peaks was performed using HOMER (v3) with a maximum motif length of 9 bp. Functional enrichment analysis of Gene Ontology (GO) annotation was conducted using the clusterProfiler R package. Enrichment significance was calculated using the hypergeometric test, with a *q*-value threshold of 0.05.

**Microscale thermophoresis (MST) assay**

The protein was labeled using the L014 Monolith NT.115 Protein Labeling Kit (NanoTemper, Munich, Germany). The Nano Temper Monolith Pico Red and Blue instrument (NanoTemper Technologies; www.nanotemper-technologies.com) was used to conduct the binding experiments. Fluorescence was measured when labeled protein and ligand were mixed and put into standard treated silicon capillaries (K022

Nano Temper, Munich, Germany). The measurements were repeated at least three times.

**Electrophoretic mobility shift assay (EMSA)**

DNA probes for the EMSA assays were generated by PCR amplification using the primer pairs listed in table. S5. The resulting promoter fragments were purified and 3′-end labeled with biotin according to the manufacturer's instructions (Thermo Fisher Scientific, Waltham, MA, USA). DNA-protein binding reactions were assembled by incubating the biotin-labeled probes with target proteins, following the EMSA protocol provided by the manufacturer (Thermo Fisher Scientific). The DNA-protein complexes were separated from unbound probes using 5% native polyacrylamide gel electrophoresis, as previously described (*2*). After electrophoresis, the gel was transferred to a membrane, and the biotin-labeled DNA was crosslinked by UV exposure. The shifted bands representing DNA-protein complexes were visualized based on biotin detection.

**Isothermal titration calorimetry analysis**

Isothermal titration calorimetry (ITC) was used to characterize the binding affinity between BDSF and EPAS1 using the ITC-200 microcalorimeter (Malvern Panalytical) according to the manufacturer's protocol. BDSF was initially dissolved at a concentration of 1 M and subsequently diluted to 0.2 mM with ultrapure water. Titrations began with a single 0.2 μL injection of 200 μM BDSF into a sample cell containing 300 μL of 20 μM protein solution, followed by 19 injections of 2 μL each. The heat changes associated with each injection were recorded, and the experiment was repeated at least three times. Data were corrected for heat changes from final injections and fitted into a one-site binding model to determine the dissociation constant ($K_D$) using MicroCal ORIGIN software (v.7).

**Molecular docking analysis**

The two-dimensional (2D) chemical structure of BDSF and EPAS1 inhibitor PT2399 were retrieved from the PubChem database. Subsequently, the three-dimensional (3D) conformation of BDSF or PT2399 was constructed and geometry-optimized using ChemBio3D. The predicted structure of EPAS1 was obtained from the AlphaFold Protein Structure Database (https://alphafold.ebi.ac.uk/entry/P97481). Prior to docking, the protein structure was prepared using the Protein Preparation Wizard in Schrödinger Suite, ensuring proper assignment of bond orders, addition of hydrogen atoms, and optimization of hydrogen-bonding networks (*3*). To define the protein's active site, the Receptor Grid Generation tool in Schrödinger was utilized, generating an enclosing grid box centered on the natural ligand to accurately encompass the binding pocket. Molecular docking of BDSF or PT2399 to EPAS1 was performed using the Glide module with standard precision (SP) settings. Post-docking, binding affinities and ligand-receptor interactions were further evaluated through molecular mechanics generalized Born surface area (MM/GBSA) calculations. Similarly, the structures of BDSF binding to other candidate proteins, including HIF-1α, HIF-3α, PSMB5, TCP1, DNAJC2, NT5C2, and EIF2S1, were predicted.

**Molecular dynamics simulation**

Molecular dynamics (MD) simulations were performed with Desmond (Schrödinger Suite 2024) to characterize particle trajectories (coordinates, velocities, energies) and evaluate the structural stability of EPAS1-ligand complexes as previously described (*4*). Systems were parameterized with the OPLS4 force field and solvated using the SPC/E water model in a cubic box; electroneutrality was achieved by adding 0.150 M NaCl. After steepest-descent energy minimization (50,000 steps), heavy-atom positional restraints were applied during NVT and NPT equilibration (50,000 steps each) at 300 K and 1 bar, followed by an unrestrained 100 ns production run. Trajectories and interactions were analyzed in Maestro 14.3. For downstream analysis of EPAS1-DNA interactions, the EPAS1 apo structure and the EPAS1-BDSF complex taken from the final 1 ns of MD, together with an AlphaFold3-predicted DNA motif, were prepared in Schrödinger (v2024): proteins were processed with the Protein Preparation Wizard (preprocessing, native ligand state regeneration, H-bond optimization, energy minimization, and water removal) and DNA with the Nucleotide Preparation Wizard. Nucleotide–protein docking was then performed using the dedicated module (70,000 ligand rotations probed, up to 30 poses returned), where lower docking scores indicate lower binding free energy and greater complex stability. Lowest-scoring poses were visualized with distinct chain coloring and surface rendering to delineate DNA binding regions for EPAS1 alone versus the EPAS1-BDSF complex.

**Sequence alignment of EPAS1 across species**

The amino acid sequences of EPAS1 from the specified species (*Mus musculus*, *Homo sapiens*, *Danio rerio*, *Rattus norvegicus*, *Gallus gallus*, *Xenopus laevis*, *Equus caballus*, *Canis lupus familiaris*, *Sus scrofa*, *Macaca nemestrina*, *Bos taurus*, *Felis catus*, and *Leopardus geoffroyi*) were retrieved from the NCBI Protein database. Multiple sequence alignment was performed using the BLASTP algorithm on the NCBI website. The resulting multiple sequence alignment was visualized using ESPript 3.0 to generate a graphical representation highlighting the conserved residues.

**Circular dichroism (CD) spectra assay**

CD analysis of EPAS1 and the EPAS1-ARNT protein complex was carried out on the Chirascan spectropolarimeter (Applied Photophysics, UK) as previously described (*5*). EPAS1 protein, EPAS1-ARNT complex, and BDSF solutions were mixed at room temperature for 30 minutes. PBS was used as the blank control. The spectra of PBS buffers were subtracted from those of the samples. The resulting spectra were analyzed to evaluate the impact of BDSF on the secondary structure of EPAS1 or

EPAS1-ARNT complex.

**Preparation of Liposome-encapsulated BDSF particles**

BDSF-loaded liposomes were prepared using the thin-film hydration method. Specifically, 2 mg cholesterol (Chol, Macklin, Shanghai, China), 2 mg DSPE-PEG2000 (Ruixi, Xi'an, China), 6 mg phosphatidylcholine (PC, MCE, Shanghai, China), and 1 mg BDSF (Abcam, Cambridge, UK) were dissolved in 5 mL chloroform within a 250 mL round-bottom flask. The organic solvent was evaporated under reduced pressure at 40 °C via rotary evaporation to form a homogeneous lipid film, followed by overnight vacuum drying to eliminate residual solvent traces. The dried film was hydrated with 1 mL PBS under vigorous shaking, subjected to bath sonication at 100 W for 10 minutes, and vortex-mixed to homogeneity. The resulting multilamellar vesicles were sequentially extruded through 400 nm, 200 nm, and 100 nm polycarbonate membranes to obtain BDSF-loaded nanoliposomes with an average particle size of approximately 90 nm. The final formulation was stored at 4 °C for subsequent use.

***In vitro* characterization of Liposome particles**

The particle size (hydrodynamic diameter) and polydispersity index (PDI) of the nanoparticles were determined using dynamic light scattering (DLS) with a Zetasizer Nano ZS instrument (Malvern Instruments Ltd., UK). The surface charge (zeta potential) was measured on the same Zetasizer Nano ZS system employing laser Doppler microelectrophoresis. The quantity of BDSF encapsulated within the liposomes was quantified using LC-MS/MS with a triple quadrupole mass spectrometer (LCMS-8060, Shimadzu Corporation, Japan) equipped with a C18 analytical column. The drug loading rate was calculated as the ratio of the encapsulated drug amount to the total initial drug input. Following drying using a Centrifugal vacuum concentrator (ZLS-2, Hunan, China), the mass of the drug-loaded liposomes was obtained, and the encapsulation efficiency was subsequently determined as the ratio of the encapsulated drug amount to the mass of the dried drug-loaded liposomes.

**Cell viability assay**

To measure the effects of BDSF, Liposome, and Liposome/BDSF particles on the cell viability of RAW264.7 cells, the cells were seeded into 96-well plates at a density of 3000 cells per well in 100 μL of DMEM and cultured overnight at 37 °C under 5% $CO_2$. The following day, the culture medium was removed, and the cells were treated with 100 μL of BDSF solution at concentrations ranging from 100 μM to 0 μM (100, 50, 25, 10, 0 μM), or with 100 μL of Liposome or Liposome/BDSF solutions at concentrations ranging from 0.5 mg/mL to 0 mg/mL (0.5, 0.25, 0.125, 0.0625, 0 mg/mL), respectively. After incubation at 37 °C for 24 hours, cell viability was assessed using the Cell Counting Kit-8 (CCK-8, Beyotime, Shanghai, China) assay, and absorbance was measured with a microplate reader (BioTek, USA).

**Bacterial infections *in vivo***

Male BALB/c mice aged 6-8 weeks were purchased from Zhuhai BesTest Biotechnology Co., Ltd. (Zhuhai, China, SCXK<Guangdong>2020-0051) and maintained by the Experimental Animals Department of Sun Yat-sen University (license SYXK<Guangdong>2024-0360, Shenzhen, China). All animal experiments were conducted in accordance with guidelines approved by Sun Yat-sen University Institutional Animal Care and Use Committee (IACUC, SYSU-2024002428). All mice were housed under specific pathogen-free (SPF) conditions with a controlled environment maintaining a temperature of 25 °C and a 12-hour light-dark cycle. All animals were provided ad libitum access to standard chow and water.

Eighty mice were divided into 8 groups, with 10 mice per group. Two groups received daily intravenous injections of either PBS or 5 mM BDSF (100 μL per injection). Investigators were blinded to treatment during outcome assessment. Two groups received daily intravenous injections of 5 mM BDSF (100 μL per injection) for four consecutive days. On day 3 of injection, mice were anesthetized with isoflurane and subsequently subjected to intratracheal instillation of either a $1\times10^7$ CFU *B. cenocepacia* strain H111 bacterial suspension or a $1\times10^7$ CFU *A. baumannii* bacterial suspension. Two groups received daily intravenous administrations of 100 μL liposome/BDSF complex (12 mg/mL total concentration, equivalent to approximately 5 mM BDSF) for four consecutive days. On day 3 of injection, mice were anesthetized with isoflurane (RWD, Shenzhen, China) and subsequently subjected to intratracheal instillation of either a $1\times10^7$ CFU strain H111 bacterial suspension or a $1\times10^7$ CFU *A. baumannii* bacterial suspension. The remaining two groups received no pretreatment. On day 3, after anesthesia with isoflurane, these mice received intratracheal administration of either a $1\times10^7$ CFU strain H111 suspension or a $1\times10^7$ CFU *A. baumannii* suspension. Lung, renal, and colonic tissues were collected for subsequent experiments. The remaining 7 mice per group were monitored daily for body weight and survival rate for a period of 8 days. Humane endpoints (e.g., > 20% weight loss, severe distress) were predefined and applied during infections.

**Colony-forming unit (CFU) determination**

The bacterial load within infected lung tissues was quantified using a standardized plating protocol. Specifically, 50 mg of lung tissue from each animal was weighed and homogenized in 1 mL of sterile PBS at 4 °C. The resulting homogenate was then subjected to three serial hundred-fold dilutions in sterile PBS. Aliquots (50 μL) of each dilution were plated onto LB agar plates. The inoculated plates were incubated at 37 °C for 24 hours. Following incubation, distinct bacterial colonies were enumerated, and the bacterial load was calculated and expressed as colony-forming units per gram of lung tissue (CFU/g).

**Single cell preparation and flow cytometric analysis**

Lung tissue specimens (two lobes per sample) were minced into fragments using sterile scalpels. The fragments were digested in 5 mL of enzymatic solution (1 mg/mL collagenase D in RPMI 1640 medium) for 1 hour at 37 °C with gentle agitation (60 rpm). Following digestion, the suspension was filtered through a 70-μm cell strainer to obtain a single-cell suspension. Cells were pelleted by centrifugation at 500 × *g* for 5 minutes at 4 °C, washed once with PBS, and resuspended in 1 mL PBS after repeated centrifugation. Cell viability and concentration were assessed using trypan blue exclusion with a hemocytometer, adjusting the final concentration to $1 \times 10^7$ cells/mL. Aliquots of 100 μL cell suspension (containing $1 \times 10^6$ cells) were incubated with 1 μL viability dye (APC-Cy7-Live/Dead (ThermoFisher, Waltham, Massachusetts, USA)) on ice for 30 minutes. Cells were then washed twice with staining buffer (RPMI 1640 containing 1% FBS), centrifuged (500 ×*g*, 5 minutes, 4 °C), and blocked with 1 mL blocking buffer (RPMI 1640 with 5% FBS) on ice for 10 minutes. A staining cocktail was prepared by adding 5 μL FITC anti-mouse CD11b (Biolegend, San Diego, California, USA), 5 μL PE Anti-Mouse CD45 Antibody (Elabscience, Wuhan, China) and 5 μL PerCP/Cyanine5.5 anti-mouse F4/80 (BioLegend, San Diego, California, USA) to 100 μL staining buffer. After centrifugation, pellets were resuspended in 100 μL staining cocktail and incubated at room temperature for 30 minutes in darkness. Post-staining, cells underwent two additional washes with staining buffer followed by resuspension in 600 μL acquisition buffer. Single-stained controls were prepared for each fluorochrome.

Flow cytometry data were acquired using calibrated instruments (Becton, Dickinson and Company, USA), with preliminary compensation adjustments performed using single-stained controls. A minimum of 50,000 events per sample were recorded. FCS 3.1 data files were analyzed in FlowJo v10.8.1, where a compensation matrix was generated automatically from single-stained controls and applied to all samples from the same experiment. Mature macrophages were quantified as the $CD11b^+F4/80^+$ population in the FITC-CD11b vs. PerCP-Cy5.5-F4/80 scatter plot, with quadrant boundaries established using fluorescence-minus-one (FMO) controls. The percentage of mature macrophages within the $CD45^+$ leukocyte population was calculated for statistical analysis.

**Histopathologic analysis**

Mice were euthanized, and lungs from each group were excised, immediately fixed in 4% paraformaldehyde overnight, paraffin-embedded, sectioned, and hematoxylin and eosin (H&E)-stained by Servicebio (Guangzhou, China) according to standard histology protocols. To assess potential toxicity of BDSF and Liposome/BDSF on kidney and colon, renal and colonic tissues were fixed in 4% paraformaldehyde overnight and processed for H&E staining as above. All stained paraffin sections were examined under light microscopy.

**RNA extraction, reverse transcription, and quantitative real-time PCR**

Total RNA was extracted from stimulated RAW264.7 cells and from renal and colonic tissues using TRIzol reagent (Invitrogen, Carlsbad, CA, USA) according to the manufacturer's instructions. Total RNA was reversely transcribed into complementary DNA (cDNA) for subsequent gene quantitative analysis. Quantitative real-time PCR (RT-qPCR) reactions were performed using SYBR Green Supermix (Promega, Madison, WI, USA) on the Applied Biosystems 7500 FAST Real Time PCR System (Waltham, MA, USA). Gene expression levels were normalized to *Actb* using specific primer pairs for mouse *Tnfa*, *Il6*, *Epas1, Map2k2*, *Map3k7*, *Map3k8, Mki67*, and *Muc2*. The PCR primers were listed in table. S5.

**Confocal microscopy**

RAW264.7 macrophages were seeded onto sterile coverslips in DMEM supplemented with 10% FBS and cultured at 37 °C until 80% confluence. Following treatment with BDSF, cells were rinsed with PBS, fixed in 4% paraformaldehyde, permeabilized with 0.1% Triton X-100, and blocked with 5% BSA in PBS for 1 hour at room temperature. Samples were incubated with anti-EPAS1/HIF-2α primary antibody (rabbit; 1:100 in blocking buffer) and labeled with Alexa Fluor 488-conjugated secondary antibody, followed by DAPI nuclear staining. Coverslips were mounted in antifade medium and imaged on a laser confocal microscope (Olympus FV3000, Tokyo, Japan).

**Enzyme-linked immunosorbent assay (ELISA)**

The supernatants from cultured RAW264.7 cells were centrifuged and stored for cytokine analysis. Quantification of cytokines was performed by using mouse TNF-α and IL-6 ELISA kits according to the manufacturers' instructions. Briefly, the capture antibodies were coated into the 96-well high-binding microplates overnight at 4 °C. After washing, the supernatants were added and incubated for 2 hours at room temperature. Followed by incubation with secondary antibodies for 1 hour and subsequent incubation with streptavidin-HRP for 1 hour at room temperature, the signals were developed by adding the TMB substrate and $H_2SO_4$ stop solution. The absorbances were obtained at 450 nm by the BioTek Microplate Reader.

**Western blot assay**

Total protein was extracted from the stimulated RAW264.7 cells by addition of lysis buffer (10 mM Tris-HCl, pH 8.0, 120 mM NaCl, 0.5% NP-40, and 1 mM EDTA) containing the protease and phosphatase inhibitors (Roche, Basel, Switzerland). Equal amounts of protein were loaded alongside a pre-stained protein ladder (1st base, Singapore) and electrophoresed in 10% SDS-PAGE gels. The resolved proteins were transferred onto the PVDF membranes (Immobilon, Millipore Corporation, Billerica, MA, USA), followed by blocking in Tris-buffered saline with Tween 20 (TBST) containing 5% milk for 1 hour at room temperature. The membranes were hybridized overnight with primary antibodies against β-Actin (#4970), phosphorylated ERK1/2 (#4377), total ERK1/2 (#4695), phosphorylated p38 (#9215), total p38 (#9212), phosphorylated JNK (#4671), or total JNK (#9252) (all 1:1000 dilution, Cell Signaling Technology, Danvers, MA, USA) and were followed by incubation with horseradish peroxidase (HRP)-conjugated anti-rabbit secondary antibody (NA934, GE Healthcare, UK) for 1 hour at room temperature. The signals were detected by the Bio-Rad chemiluminescence system (Bio-Rad, Hercules, CA, USA) with Amersham Hyperfilm ECL reagent (GE Healthcare, UK), SignalFire™ ECL reagent (Cell Signaling Technology), or SuperSignal™ West Atto Ultimate Sensitivity Substrate (Thermo Fisher Scientific).
